# Impact of Norway spruce pre-degradation stages induced by *G. trabeum* on fungal and bacterial communities

**DOI:** 10.1101/2021.11.02.466070

**Authors:** Nicolas Valette, Arnaud Legout, Barry Goodell, Gry Alfredsen, Lucas Auer, Eric Gelhaye, Delphine Derrien

## Abstract

In forests ecosystems, fungi and bacteria are key actors in wood degradation. However, few studies have focused on the impact of fungal pre-degradationon bacterial and fungal communities. Coniferous forests are dominated by Brown rot fungi that are involved in earliest phase of lignocellulose breakdown, and therefore, influencing the second phase of microbial colonization. This study aimed to investigate the first microbial species colonizing after intermediate and advanced pre-degradation by the brown rot fungus *Gloeophyllum trabeum*. Using Illumina metabarcoding, bacterial and fungal communities were monitored after 70 days from *Picea abies* wood blocks placed between forest floor and topsoil. Chemical composition of the wood samples was determined for each of the pre-degradation stages. We observed significant changes in the bacterial and fungal communities associated with the pre-degradation of *P. abies*, and that wood substrate condition acted as a gatekeeper for both microbial communities. Our data also suggested that fungal and bacterial communities could interact and act in a synergistic way during the wood decomposition process.

## Introduction

Forests covers ~30% of global terrestrial area but forest soils account for more than 50% of the total soil organic C stocks (Carvalhais et al., 2014). In forest ecosystems, coarse woody debris is the primary source of biomass for soil organic matter (SOM) formation. It is composed primarily of polysaccharides and lignin at 65-80% and 20-35% (Mao et al., 2019) respectively. The wood-rotting Basidiomycota fungi play a central role in woody litter degradation. They are the major players able to remove or circumvent the lignin barrier that hinders access to plant polysaccharides, which are the major plant components that can support microbial growth (Arantes & Goodell, 2014). Although it is recognized that there is intergradation in the different types of wood decay which can occur (Riley et al., 2014), the decay fungi are typically categorized into two distinct but related functional groups of Basidiomycota: the brown rot and white rot fungi (Floudas *et al*., 2012). Indeed, the emergence of the brown rot lineage is reported to be the result of multiple evolutionary events stemming from white rot ancestors with the contraction of gene families involved in lignin decomposition (Floudas *et al*.,2012; Nagy et al., 2017). White rot fungi are sub-divided into species able to degrade cellulose, hemicelluloses and lignin simultaneously, and species which selectively depolymerize lignin before polysaccharide part, primarily using enzymatic systems. Brown rot fungi employ a different biodegradative strategy and they can remove the carbohydrate from plant tissues by disrupting and then leaving a residue of modified lignin. The brown rot fungi generate hydroxyl radicals via what has been described as a chelator-mediated Fenton (CMF) reaction (Goodell et al., 2020). These hydroxyl radicals depolymerize lignin, which then allows polysaccharides to diffuse out of the cell wall where fungal enzymes would primarily be located (Goodell et al. 2017; Zhu et al., 2020) and may also provide greater accessibility of enzymes to the wood cell wall depending on what species of wood and fungi are involved (Arantes & Goodell, 2014; Goodell et al., 2017; Presley et al., 2018).

The order and timing of species colonization of wood residues during community assembly can affect species abundances and functional diversity, affecting wood decay rate (Fukami *et al*., 2015Zhang et al., 2016; Zhang et al., 2019)). Relative to the decay of wood in soils, the ability of the microbial community to establish, and how this is impacted by the intrinsic properties of the ecosystem, is also very important relative to the successional establishment of primary and secondary microbial species (Zhalnina *et al*., 2014; Lopez-Mondejar *et al*., 2015; Dickie *et al.,* 2012; Hiscox *et al*., 2016). In living trees, the presence of wood-decay fungi has been observed, meaning that deadwood can already be well colonized by fungi before it falls to the forest floor (Boddy *et al*., 2001). In contrast, Johnston *et al*. (2016) suggested that deadwood bacteria are not endophytes but establish primarily from soil contact, and would therefore be selected by associated primary fungi. The importance of the primary fungal colonizer is underlined by several field experiments where the presence of different decay fungi in wood both filtered and regulated the establishment of secondary microbial communities (Hiscox *et al*., 2015; Hiscox *et al*., 2016; Jonhston *et al*., 2019; Christofides *et al*., 2019).

Recently, Viotti *et al*. (2021) highlighted the different bacterial and fungal community structure dynamics occurring between oak sapwood and heartwood, suggesting that wood physiochemical properties are important in the microbial assemblage history. Indeed, wood decay rates were correlated with the chemical composition of wood, and decay was strongly reduced both by lignin and extractives content (Kahl et al., 2017). Heartwood extractives are known to have a composition and level of antifungal activity that varies widely within tree species (Valette et al., 2017). This variability influences the colonization of wood by fungi that must detoxify these molecules (Perrot et al., 2020). Consequently, wood physiochemical properties are important drivers of microbial wood colonization and influence, especially the establishment of the pioneer organisms in succession. Thus, during wood degradation processes, both fungi and bacteria modify wood physiochemical properties, and particularly in heartwood, they must first employ their secreted metabolites to cope with extractive molecules and then follow this by enabling cellular machinery which can degrade major wood cell wall polymers.

While some studies have already addressed the influence of primary colonizing fungi on bacterial or fungal colonization strategies, little attention has been paid to examining joint fungal-bacterial colonization and succession. Most studies have focused on the primary colonizing organisms and not on the chemical properties of the decaying wood substrate (Folman et al., 2008; Hervé et al., 2014; Hiscox *et al*., 2016; Jonhston *et al*., 2019; Christofides *et al*., 2019). To help fill this knowledge gap, we carried out a soil incubation using Norway spruce (*Picea abies*) blocks with pure cultures of the brown rot fungus *Gloeophyllum trabeum* to produce spruce blocks decayed to different degrees. This aim to reproduce the effect of fungal community dominated by brown rot fungi during the early stages of wood decay in coniferous forest (Ottosson et al., 2013).The spruce blocks were then transferred to unsterile soil; buried in the forest floor of a Norway spruce forest in NE France and incubated for 70 days. We hypothesized that both fungal and bacterial secondary colonizers would attack the wood differently depending on the stage of wood decay introduced to the forest soil. To assess this, fungal and bacterial communities in the blocks were monitored over time using high-throughput amplicon sequencing. We also investigated whether the secondary decomposers originated from the forest floor or from the litter, and observed putative fungi/bacteria interactions using coincident data pattern assessment.

## Results

### Chemical composition of Norway spruce pre-degraded by Gloeophyllum trabeum prior in situ incubation

Laboratory decay (pre-degradation) of Norway spruce by the brown rot fungus *G. trabeum* induced substrate mass loss and changes in wood biochemistry (Table S1; Fig. 1). Mass loss increased with pre-degradation duration, but was quite variable (Fig. 1A). For example, 8.1 ± 1.9, 15.2 ± 8.4 and 24.4 ± 4.7% of the initial wood mass was lost after 40, 105 and 165 days of pre-degradation decay, respectively. Pre-degradation by *G. trabeum* also resulted in a relative decrease in polysaccharides, with a relative increase in both the soluble components and in lignin; which is in accordance with the mechanisms employed by brown rot fungi (Green & Highley, 1997; Arantes & Goodell, 2014; Bader *et al*., 2012). Wood blocks were grouped according to their mass loss (independent of their preincubation time) as either: non-decayed wood (0%, n= 6), intermediate decayed wood (13.36 ± 0.17%, n = 6) or advanced decayed wood (31.71 ± 1.07%, n = 6) (Table S1; Fig. 1B).

**Fig. 1:**
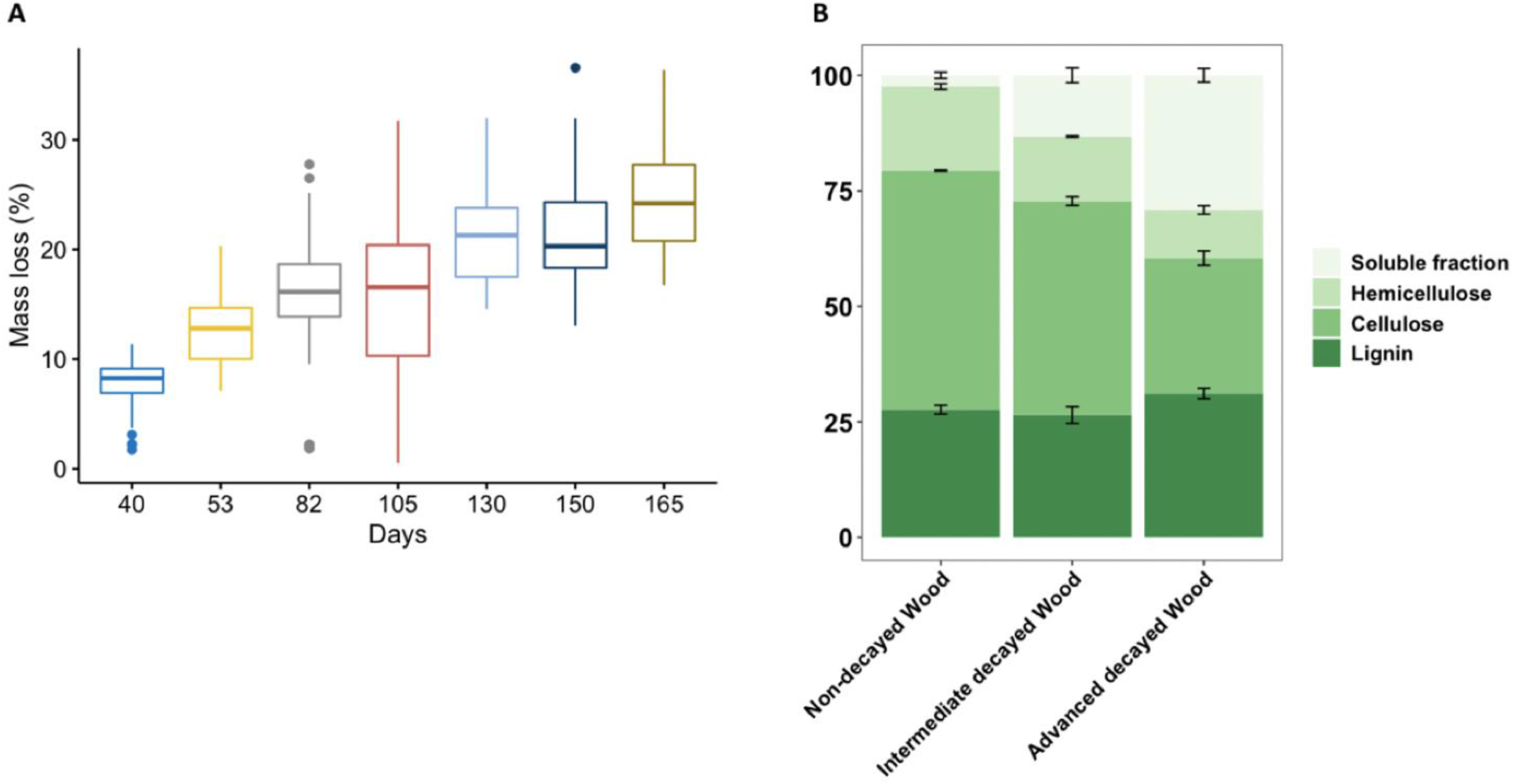
A) Norway spruce mass loss variability in laboratory pre-decayed wood induced by *G. trabeum* and B) Chemical composition of the laboratory pre-decayed samples when grouped as either non-decayed, intermediate decayed, or advanced decayed blocks based on mass loss.

### Norway spruce decay stages decreased microbial diversity and shaped the structure of microbial communities

After 70 days of burial in litter on the forest floor, no additional mass loss was observed in any of the three wood groups beyond the mass loss observed during the laboratory pre-decay period.

The diversity analysis of the bacteria colonizing the pre-degraded wood blocks based on 31 samples was successfully sequenced and provided 1,204,722 high quality reads (554,534 sequences for the incubated wood blocks, 323,776 sequences for the forest floor, and 288,031 sequences for the forest soil). Colonizing fungi diversity analysis was based on 29 samples, which provided 1,201,401 high quality reads (575,581 sequences for the incubated wood blocks, 356,480 sequences for the forest floor and 269,340 sequences for the forest soil). Sequences were rarified at 3,488 and 2,886 per sample and distributed into 582 OTUs for bacteria and 526 for fungi. Alpha microbial diversity was assessed using the Chao1 richness estimator, Shannon’s diversity index and Pielou’s evenness index and all were significantly influenced by the wood, forest floor or soil environment (Fig. 2A and 2B). Wood environments significantly reduced bacterial and fungal richness in comparison to soil and forest floor, independent of the pre-degradation stages. Moreover, pre-degradation of Norway spruce by *G. trabeum* induced a decrease in both bacterial and fungal diversity compared to the other environments (Fig. 2A and 2B).

**Fig. 2:**
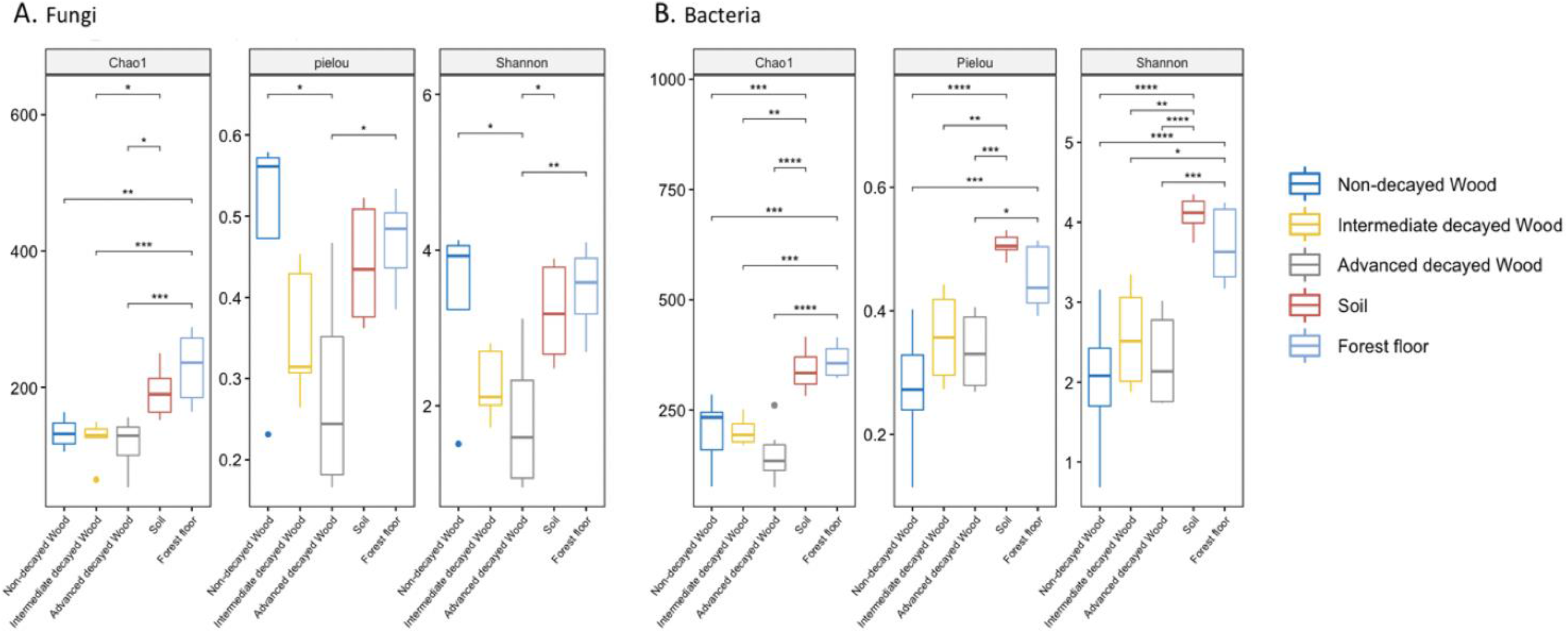
Microbial diversity analysis measured for the Non-decayed, Intermediate-decayed, Advanced-decayed woods, and also Soil and Forest floor using calculated Chaol richness estimator, Pielou’s evenness index and Shannon’s diversity index for (A) fungi and (B) bacteria. The influence of the environments was tested using a pairwise comparison (p values adjusted using the Holm method).

Bacterial and fungal community structure was strongly influenced by the substrate environment (Fig. 3). Indeed, the change of ecological niche explains 48% of the variance in the OTU composition respectively for bacteria (Fig. 3A) and 51% for fungi (Fig. 3B). PERMANOVA analysis confirmed that wood environments (non-decayed wood, intermediate and advanced decay wood) significantly impacted bacterial (*R*^*2*^ = 0.27, *P* < 0.001) and fungal (*R*^*2*^ = 0.35, *P* < 0.001) communities compared to the soil and forest floor. Inside wood environments, pre-degradation of Norway spruce explained around 32% of the variance in OTU composition for bacteria and fungi. Although, pre-degradation of Norway spruce by *G. trabeum* influenced the overall structure of both bacterial and fungal diversity compared to the other environments, no major difference in microbial community structure was observed between intermediate and advanced wood decay. This indicates that initial colonization of the pre-degraded blocks, once moved to the forest floor, thus appears to play an important role in maintaining an established colony. It should be noted that the bulk of *G. trabeum* mycelium from the pre-degradation phase would have been killed in the drying phase of the blocks prior to installation at the forest site.

**FIG 3.**
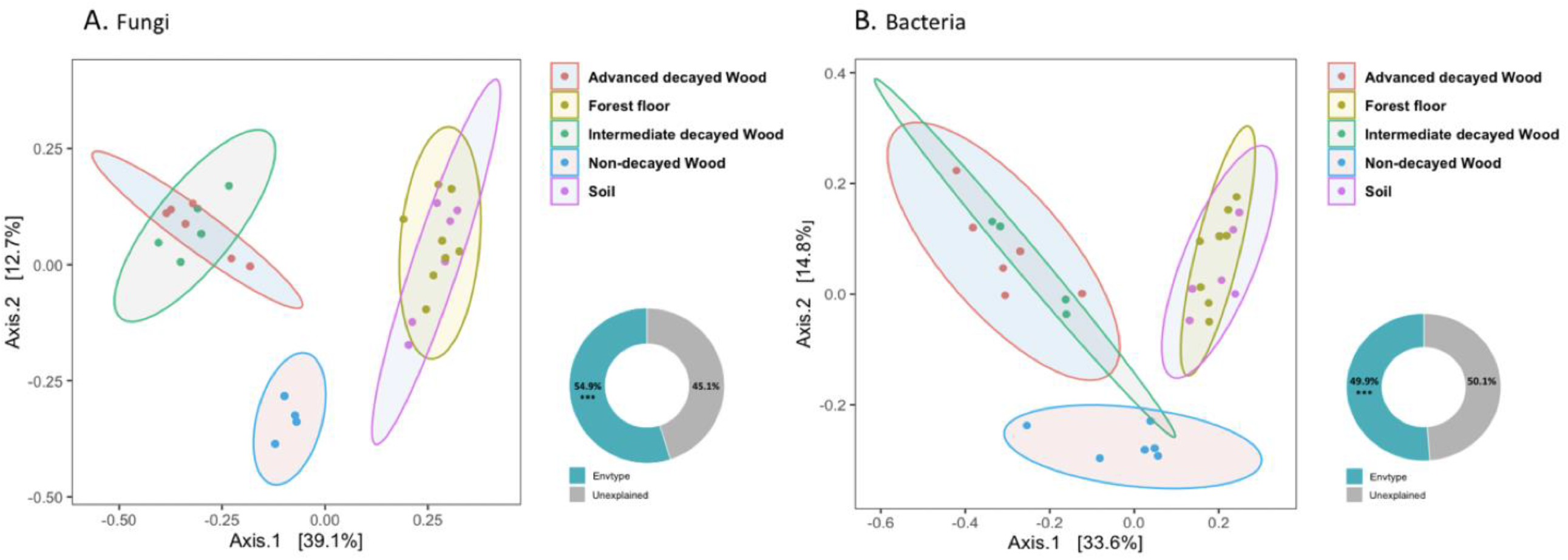
Principal coordinates analysis (PCoA) of fungal (A) and bacteria; (B) communities found in different environments The PCoA plots were based on fungal and bacterial OTUs Unifrac distance matrices. The donut charts represent the variances (%) explained or not based on permutational multivariate analysis of variance (PERMANOVA); *** P < 0.001.

### Bacteria and fungi taxonomic composition in pre-degraded Norway spruce and forest litter differed from that in the forest soil

Differences in fungal and bacterial taxonomic composition highlighted the strong influence of environment on relative microbial abundance (Fig. 4A and B).

**Fig 4.**
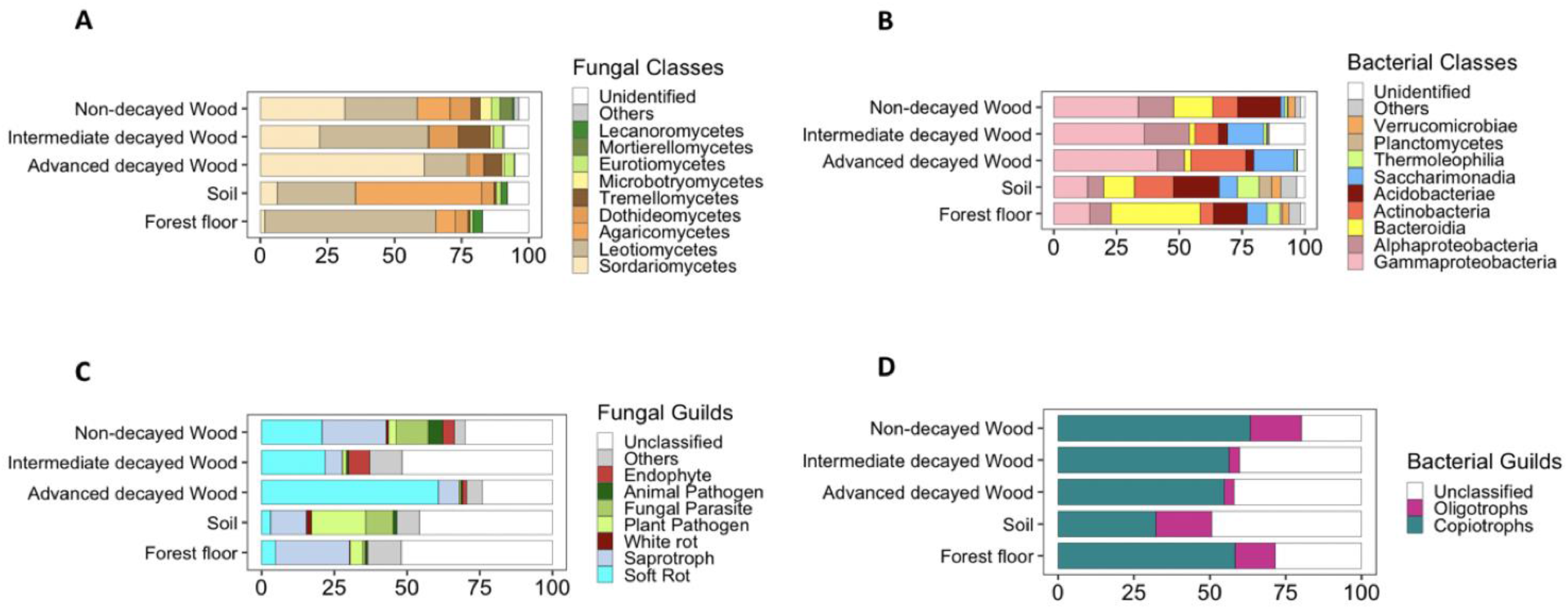
Relative abundance of (A) fungal and (B) bacterial classes and relative abundance of ecological guids for (C) fungi and (D) bacteria according to environment types.

Fungal Phyla were dominated by Ascomycota in non-decayed wood, intermediate decay wood, advanced decay wood and in the forest floor. Conversely, Ascomycota and Basidiomycota were almost equally distributed in soil. Interestingly, more Ascomycota were present in non-decayed and intermediate pre-degraded wood compared to soil and forest litter, and Ascomycota increased further still in advanced pre-degraded wood (Fig. 4C). The changes were also observed at the class level: the forest floor was dominated by *Leotiomycetes* (65%), soil was primarily colonized by *Agaricomycetes* (48%) and *Leotiomycetes* (29%), whereas wood environments with non-decayed wood, intermediate decay wood and advanced decay wood were settled by *Sordariomycetes* (40 ± 21%) and *Leotiomycetes* (29 ± 14%) (Fig. 4A). The classes with a lower abundance were also influenced by the environment since all wood types were enriched by *Tremellomycetes*, *Eurotiomycetes* and *Microbotryomycetes* classes in comparison to forest floor and soil (Table S2). Interestingly, a potential shift in Basidiomycota classes was observed between *Agaricomycetes* in non-decayed wood to *Tremellomycetes* in both intermediate decay and advanced decay wood (Fig. 4A). The Tremellomycetes are characterized as fungal parasites and their presence in the pre-degraded samples is consistent with this fungal lifestyle. Non-decayed wood was the only environment with *Mortierellomycetes* (Fig. 4A). At the Genus level, wood environments were enriched in several genera such as *Penicillium*, *Scleroconidioma*, *Cadophora*, *Krasilnikovozyma*, *Oidiodendron* or *Kendrickiella* compared to the soil and forest floor sampling. The domination of the *Trichoderma* genus in non-decayed wood was reinforced in the intermediate and advanced pre-degraded wood showing the influence of the pre-degradation. Moreover, intermediate decay and advanced pre-degraded wood showed an enrichment in *Saitozyma* and an impoverishment in *Tylospora* and *Mortierella* in comparison to non-decayed wood (Table S2).

The relative abundance of bacterial classes was dramatically different comparing the pre-degraded wood blocks to the forest soil and litter (Fig. 4B). The forest floor was dominated by the *Bacteroidia* class (35%) whereas the major class in soil were *Acidobacteriae* (18%), *Actinobacteria* (15%), *Gammaproteobacteria* (13%), and *Bacteroidia* (12%). Wood environments, independently of the decay stages, were populated by a high proportion of *Alphaproteobacteria* (14%) and *Gammaproteobacteria* (37%). The influence of the pre-degradation was also observed on less abundant classes with a decrease in the relative abundance of *Bacteroidia* and *Acidobacteriae* and an increase of *Actinobacteria* and *Saccharimonadia* observed in pre-degraded wood in comparison to control wood (Fig. 4B). The influence of these different environments was also found at the family level (Table S2). The forest floor contained the largest amount of *Sphingobacteriaceae* (27%), with other environments containing less of this family. Wood environments were dominated by several other families such as *Burkholderiaceae* and *Microbacteriaceae* in comparison to soil and litter. Bacterial abundance was also influenced by the pre-degradation stage with *Burkholderiaceae* over-represented in the intermediate- (34%) and advanced- (39%) pre-degraded wood compared to non-decayed control wood (9%), whereas *Microbacteriaceae* was found in greater proportion in the intermediate- (7%) and advanced- (19%) pre-degraded wood compared to non-decayed control wood (1%) (Table S2). Moreover, wood pre-degradation also positively influenced the abundance of other families such as and *Acetobacteraceae*, and reduced the presence of *Sphingobacteriaceae*, *Acidobacteriaceae* and *Sphingomonadaceae* families (Table S2).

### Bacterial and fungal guild composition

Despite many OTUs that could not be assigned to any guild, a higher proportion of copiotroph bacteria compared to oligotroph bacteria, were observed in all environments (Fig. 4D). Although, the percentages of copiotrophic bacteria were equivalent (around 54 - 64%) when comparing forest floor, non-decayed wood, intermediate decay wood and advanced decay pre-degraded wood, the soil showed a lower proportion of copiotrophs with only 32% (Fig. 4D). The increased number of copiotropic bacteria corresponded to a lower relative abundance of *Alphaproteobacteria*, *Gammaproteobacteria* and *Bacteroidia* in soil than in other environments (Fig. 4B). Moreover, the relative abundance of oligotrophic bacteria in intermediate and advanced pre-degraded wood (around 3%) was reduced compare to the forest floor, soil, and non-decayed wood (respectively ranging from 13 to 18%) (Fig. 4D). This could be explained by a significant decrease in *Acidobacteriae* in intermediate decay wood and in advanced decay pre-degraded wood (Fig. 4B). Interestingly, the type of environment seemed to play a role in the bacterial guild composition with soil having strongly reduced copiotrophic bacteria populations whereas intermediate and advanced pre-degraded wood having proportionally decreased oligotrophic bacteria.

The analysis of fungal guilds highlights the different ecological compositions of both environments (Fig. 4C), even given that up to 50% of the OTUs were unassigned in the soil, forest floor and intermediate decay wood environments. Among the assigned OTUs, the forest floor was dominated by saprotrophs (25%), the soil population consisted primarily of plant pathogens (19%) and saprotrophs (17%), wood environments with non-decayed wood, intermediate decay wood and advanced decay wood were represented by saprotrophs at 44%, 28% and 68% respectively. This ecological guild was dominated by soft rot at 21%, 22% and 61% respectively for non-decayed wood, intermediate decay wood and advanced decay wood (Fig. 4C). All wood types showed a higher proportion of soft rot when compared to the soil and forest floor, and advanced wood decay seemed to strongly favor colonization by this fungal guild. This phenomenon corresponded to an increase in the relative abundance of *Sordariomycetes* (Fig. 4A). An important variation was also observed for saprotrophic organisms, with their presence being reduced in the intermediate (6%), and advanced (7%) pre-degradation blocks in comparison to non-decayed samples (22%). Endophyte fungi were only observed in wood environments and confirmed the impact of wood on fungal guild composition (Fig. 4C).

### Abundant fungi and bacteria in wood co-occurred in all environments

Analysis of the most abundant OTUs from the forest floor, soil, non-decayed wood, intermediate decay wood and advanced decay wood showed that 116 bacterial and 128 fungal OTUs displayed at least 1% abundance in at least one sample. Among the most abundant OTUs, 11 fungal and 30 bacterial OTUs were selected because of their specificity to intermediate and advanced pre-degraded wood samples (Fig. 5B). Their relative abundance in all environments were used in a Spearman rank correlation at the family and species level respectively for bacteria and fungi (Fig. 5). This analysis highlighted a positive correlation (p <0.001) between bacterial species belonging to three families: *Burkholderiaceae*, *Microbacteriaceae* and *Acetobacteraceae*, and several fungal species such as *Trichoderma spirale*, *Penicillium aethiopicum*, *Saitozyma podzolica* and *Scleroconidioma sphagnicola* (Fig. 5A). These results suggest a potential interaction between fungi and bacteria, potentially which may have occurred as part of the wood degradation process. Moreover, when positive correlations between organisms were observed (in all environments) this potentially indicated that specific and potentially positive interactions occurred for the species involved.

**Fig 5.**
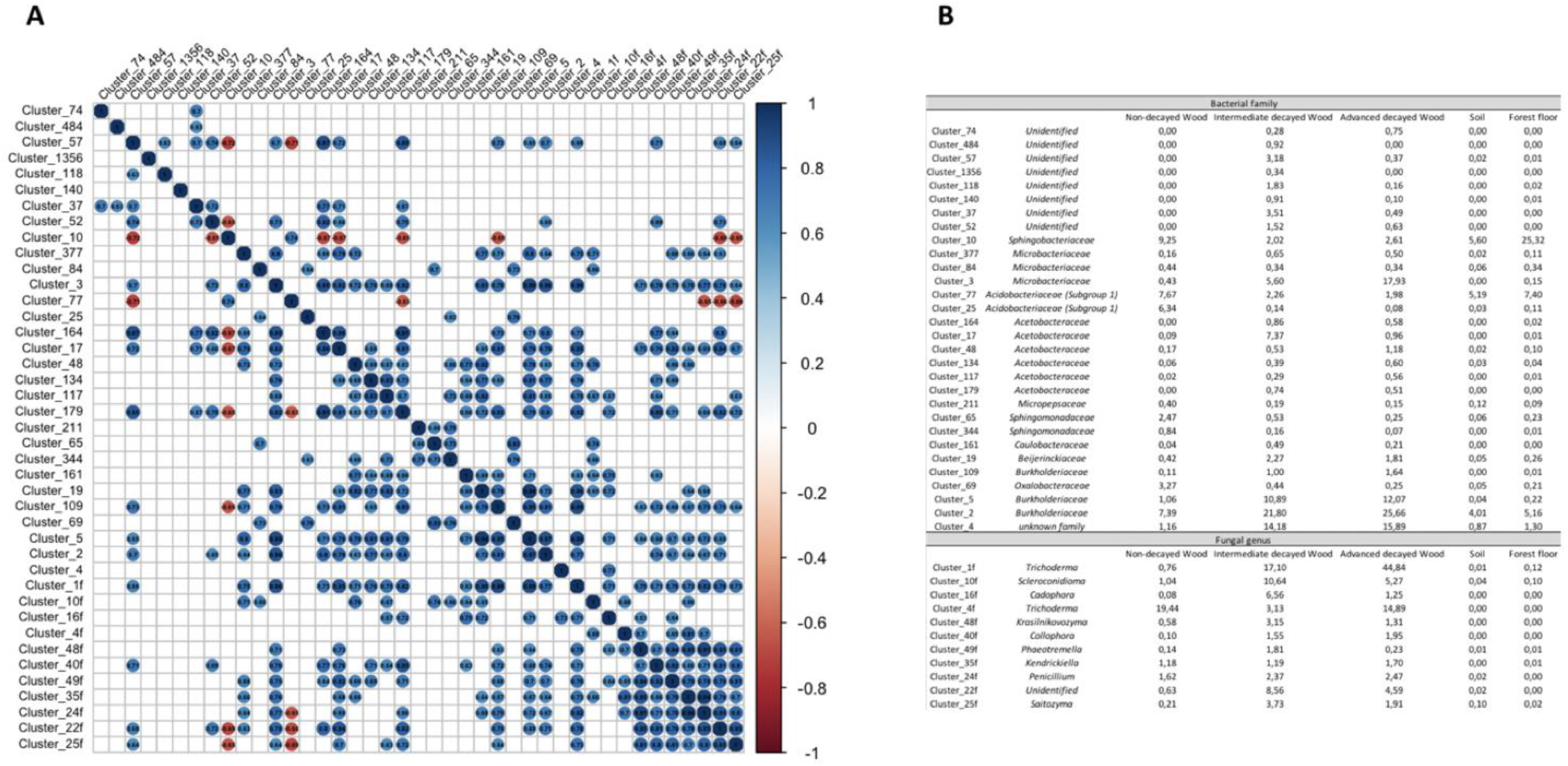
Spearman rank correlation based on (A) fungal and bacteria OTUs, and (B) abundance or specificity to wood environments. Non-significant correlations (p > 0.001) are represented by white squares (5A).

## Discussion

In order to study the beginning of the second phase of microbial colonization, we reproduced the early stages of wood decay from coniferous forest with the Norway spruce pre-degradation induced by a brown rot fungus. We demonstrated that the pre-degradation wood decay stages promoted selection of bacterial and fungal communities after samples were transferred to forest floor and forest litter environments. Significant differences in bacterial and fungal communities occurred and these were influenced by both the pre-degradation stage as well as the forest environment. We observed significant differences between wood microbial communities compared to soil and litter, suggesting an important selection pressure by wood.

### Pre-degradation of Norway spruce by G. trabeum enhanced selection of competitive fungal species

Our results demonstrated the influence of Norway spruce pre-degradation by *G. trabeum* on fungal secondary colonizers. The *Trichoderma* genus dominated in all wood types, with its proportion increasing as pre-degradation increased to represent around 60 % of total species abundance in the advanced pre-degraded wood samples. *Trichoderma* spp. on Norway spruce environments is well known for its ability to secrete a variety of CAZymes involved in cellulose degradation (Strakowska *et al*., 2014; Blaszczyk *et al*., 2016; Krogh *et al*., 2004; Isodorov *et al*., 2016). *Trichoderma* spp. are also well described to outcompete other microorganisms through the secretion of antifungal compounds, and to use dead mycelia as their carbon-nutrient source (Hiscox *et al*., 2018; Schuster & Schmoll, 2010; Gómez-Brandón *et al*., 2020). Moreover, a shift was observed in Basidiomycota populations from *Agaricomycetes* in the non-decayed wood to *Tremellomycetes* in intermediate and advanced pre-degraded wood. Species from *Agaricomycetes* are described as saprotrophic and as they are involved in wood decay processes (Nguyen *et al*., 2016). Conversely, species from the *Tremellomycetes* are described as mycoparasites and possess genes allowing them to recycle carbon from dead mycelia and degraded wood polysaccharides (Kurtzman & Boekhout, 2017; Oberwinkler & Lowy, 1981; Kim *et al*., 2018). Others dominant genera observed in intermediate and advanced decayed wood, included *Penicillium* and *Scleroconidioma,* which are known to produce CAZymes involved in some types of cellulose degradation (Koukol & Kovářová, 2007; Krogh *et al*., 2004). Overall, the pre-degradation of wood by *G. trabeum* drove microbial selection toward fungal species involved in lignocellulose degradation and also toward species that could outcompete other organisms, and that utilize live and dead mycelium from other species as a nutrient source. Therefore, depolymerization of the wood cell wall during the pre-degradation phase, as well as the presence of dead mycelium from *G. trabeum,* provided a favorable environment for the establishment and development of these fungal species.

### Potential roles of bacteria selected by the pre-degradation of Norway spruce

Bacterial colonization of the intermediate and advanced pre-degraded samples resulted in a decrease of oligotrophic bacteria corresponding with a lower proportion of *Acidobacteria*. This lead to an increase of the *Proteobacteria* : *Acidobacteria* (P/A) ratio suggesting that the high nutrient content, and especially carbon, of the pre-degraded wood aided in this selection (Kielak *et al*., 2016).

*Burkholderiacea*e was the most important family in pre-degraded wood. Although this family possesses many genes involved in lignocellulosic degradation, Johnston *et al*. (2016) reported numerous associations of *Burkholderiaceae* with fungi suggesting another role for this family in wood decay. Indeed, many species are known to be N2 fixing bacteria and can provide nitrogen essential for fungi (Gómez-Brandón *et al*., 2020). Moreover, they could also be involved in carbon cycling of lignin breakdown products from fungal activity. They are known to be able to degrade mono-, poly- and heterocyclic aromatic compounds (Morya *et al*., 2014; Jung *et al*., 2015; Perez-Pantoja *et al*., 2012). Others bacterial families were also favored by Norway spruce after pre-degradation, notably species in the *Microbacteriaceae* family belonging to the *Actinobacteria*. Interestingly, Viotti *et al*. (2021) suggested that this family could also be involved in carbon recycling from dead mycelia. Moreover, many of these bacteria are saprophytic and chemoheterotrophic, which allows them to use a wide variety of nutritional sources ranging from polysaccharides to lignin-related aromatic compounds and polymeric lignin, potentially suggesting direct degradation of lignocellulosic material (Barka *et al*., 2016; Schellenberger *et al*., 2009; Zimmerman *et al*., 1990; Wang *et al*., 2016). Other *Actinobacteria* have been described as endophytic diazotrophs in Norway spruce logs and could potentially provide nitrogen for fungal activity (Puri *et al*., 2018).

Additional bacterial families were also enhanced in pre-degraded wood. The *Acetobacteraceae* family increased in both intermediate- and advanced- pre-degraded wood. This family is known for their capacity to fix nitrogen and for their ability to solubilize phosphorous and zinc (Suman *et al*., 2001; Saravanan *et al*., 2007). In wood decomposition processes, access to both nitrogen and phosphorous promoted by bacteria could favor fungal activity. We also observed an increase in species from the *Saccharimonadia* family belonging to the *Patescibacteria* phyla. The small size of this family’s genomes as well as the description of the epibiotic parasitic lifestyle of the strain TM7x (CPR/*Patescibacteria*) suggests that the *Saccharimonadia* may have a symbiotic lifestyle (Lemos *et al*., 2020; He *et al*., 2015). Prior research reinforces this concept as a co-occurrence analysis also suggests that *Patescibcateria* could interact with members affiliated with *Bacteroidia*, *Alphaproteobacteria*, *Gammaproteobacteria* and *Actinobacteria* species which are abundant in pre-degraded Norway spruce (Lemos *et al*., 2019).

Overall, the data suggest that the predominant bacteria that established in pre-degraded wood were able to metabolize compounds from the pre-degraded wood, and they also were also able to provide nutrients to the fungi to promote fungal growth. In early wood decay stages, bacteria were not directly involved in lignocellulose degradation but our data suggest that they established in support of fungal activity.

### Bacteria and fungi co-occurred in environments under mutualistic interaction

The abundance of several bacterial families (*Burkholderiaceaea*, *Microbacteriaceae* and *Acetobacteraceae*) showed a positive correlation with two fungal genera (*Penicillium* and *Trichoderma*) in all environments. In the context of wood degradation, bacteria could potentially take up carbon from fungal hyphae exudates or lignocellulose degradation products, while at the same time also supplying the fungi with essential nutrients such as nitrogen, phosphorus or zinc (Zhang *et al*., 2020; Saravanan *et al*., 2007; Schellenberger *et al*., 2009; Kirker et al., 2017; Zelinka et al., 2021). Moreover, some bacteria can detoxify antifungal compounds and therefor they may also help to synergistically protect fungal species in the same environment (Nazir *et al*., 2014). In all environments, the co-occurrence of bacteria and fungi may promote the growth and distribution of bacteria. Indeed, in soil, fungal hyphae form a belowground network that specific bacterium have been reported to navigate subject to localized environmental conditions, to promote their migration (Yang Pu 2017). Soil is a sporadic source of nutrients that are often concentrated in hotspots and ephemerally available. The use of this semi-continuous network by bacteria would enhance their ability to discover new nutrient-rich environments. Bacterial dispersal along fungal hyphae could be the result of mutualistic interactions, with soil inhabiting fungi being known to mobilize pollutant-degrading bacteria with resultant mediation of bacterial diffusion to the polluted sites, thereby enhancing soil bioremediation (Kohlmeier *et al*., 2005; Furuno *et al*., 2010). Finally, bacteria and fungi co-exist and interact in a variety of ways to impact their micro-environments in ways that will release nutrients and promote synergistic growth.

## Conclusion

The pre-degradation of wood by *G. trabeum* causes an important modification of both bacterial and fungal communities. These communities interact synergistically, i.e, fungi are the main decomposers of organic matter to provide carbon (wood degradation products and fungal hyphal residues and exudates) for bacteria. In-turn, the bacteria provide nutrients (nitrogen, zinc or phosphorous) to the fungi, and protect fungi against antagonistic compounds produced both by other bacteria and invading fungi. In our research, particularly with pre-degraded wood, wood environments promoted a strong decrease in fungal and bacterial communities compared to soil and forest floor environments.

## Experimental procedures

### In vitro and field site degradation

Norway spruce blocks (Picea abies Karst) (50×25×15 mm) originating from Norwegian forest stands were placed in petri dishes containing malt agar medium and four 7mm inoculum plugs of *Gloeophyllum trabeum* (strain 88913) for decay according to EN 113. Cultures were incubated at 25°C for 50, 80, 105, 130, 150 and 165 days and wood mass loss on 3 replicate sacrificial samples was then measured on the dried samples (60°C, 12 h). After laboratory decay exposure (pre-degradation), the wood samples were installed in Spring 2019 at the Breuil-Chenue experimental forest site, in the Morvan area in France (47°18′N, 4°5′E). The forest is located on a plateau at 640m, the annual rainfall is 1280 mm and the average temperature is 9°C. The soil is an alocrisol that developed over the ‘La Pierre qui Vire’ leucogranite with a topsoil pH ranging between 3.8 and 4.5 (Baize & Girard, 1988; Levrel & Ranger, 2004)(Table S3). In 1976, a part of the native forest was clear-cut and the area was planted with several mono-specific plantations (1000 m2 for each). Among these, one plot that was planted with Norway spruce was selected for this study. We selected a 90 x 45 cm area and carefully removed the forest floor litter layer for placement of bagged wood block samples in a block design on the topsoil as described below. Polyamide bags with a fabric mesh size of 1 mm were used to each contain a single wood block. Six replicates were used for each treatment including the undecayed control. Treatments included “undecayed controls” (0% laboratory mass loss), “intermediate” decayed samples (13.36 ± 0.17% laboratory mass loss), and “advanced” decay samples (31.71 ± 1.07% laboratory mass loss). The bagged blocks were uniformly distributed throughout the area, covered with the original forest floor litter layer, and then after 70 days these samples were collected. Eight forest floor litter layer samples, and eight soil cores from the 0-5 cm mineral horizon were also collected (Fig. S1). All samples were transported to the laboratory frozen in ice, and then were stored at −80°C. Before DNA extraction, samples were comminuted using a cryomill (Retsch GmbH, Germany) following the manufacturer’s instructions and three subsamples (700 mg) of soil, litter or wood were used for DNA extraction.

### Wood biochemistry of Norway spruce

A modified Van Soest protocol with “Fibrebag” technology (C. Gerhardt GmbH & Co., Germany) (Van Soest & Wine, 1967; Van Soest & McQueen, 1973) was used to filter the solubilized wood polymers during each sequential solvent treatment stage. Briefly summarized, these stages were: 1) Extractives removed first through solubilization using a neutral detergent fiber solution (sodium dodecyl sulfate, sodium EDTA, sodium phosphate monobasic, sodium tetraborate decahydrate and triethylene glycol, with pH ranging between 6.9 and 7.1). 2) Hemicellulose was then solubilized using an acid detergent fiber solution (CTAB dissolved in 1l of sulfuric acid 0.5 mol/l). 3) Finally, an acid detergent treatment (sulfuric acid, 72 %) was used to solubilize cellulose, leaving lignin as final product. Each solubilized fraction was collected and dried overnight at 105°C before weighing to determine mass loss as compared to undecayed reference samples.

### DNA extraction, PCR amplification and sequencing

Genomic DNA was extracted from 700 mg of comminuted materials (soil, forest floor litter or wood) using a RNeasy powersoil DNA elution kit following the manufacturer’s instructions (Qiagen GmbH, Hilden, Germany). No DNA was retrieved from the non-decayed wood samples, so a second “surface extraction” method was then used with the same kit assuming that, in these samples, any microbial colonization would have occurred mainly at the surface of the wood blocks. Using this method, DNA was extracted from surface-etched non-decayed wood blocks with a success rate of 75%. Amplification of the fungal ITS2 region was then performed using forward primer ITS86f (5’-GTGAATCATCGAATCTTTGAA-3’) and reverse primer ITS4r (5’-TCCTCCGCTTATTGATAGTC-3’)(Op De beeck *et al*., 2014). For bacterial communities, the 16S rRNA V4 region was amplified with forward primer 515f (5’-GTGCCAGCMGCCGCGGTAA-3’) and reverse primer 806r (5’-GGACTACHVGGTWTCTAAT-3’) (Caporaso *et al*., 2011). For each DNA extraction, three independent PCR reactions were carried out in a final volume of 50 µl with Herculase II fusion DNA polymerase (Agilent), then pooled and confirmed on gel electrophoresis for both fungi and bacteria. The PCR conditions used were 95°C for 3min, 33 cycles of 45 s at 95°C (denaturation), 55°C for 30 s (annealing) and 72°C for 30 s (extension), followed by 10 min 72°C. The multiplexed and the Illumina MiSeq sequencing were performed by Genoscreen Society (Lille, France, https://www.genoscreen.fr/fr/). The raw MiSeq sequences were submitted to the Sequence Read Archive (SRA) and are available under the BioProject ID: PRJNA775075.

### Miseq sequences processing and bioinformatic analyses

Sequences were demultiplexed with CASAVA v1.0 software (script PERL ConfigureBclToFastq.pl) and merged with a FLASH tool configured to suppress the PCR primer (100% of identity), to trim extremities with a score ≤ Q30, and to maintain a minimal length of 30 nucleotides with an identity yield of 97 % (Magoc & Salzberg, 2011). The paired sequences (read 1 and read 2 merged) were then processed with FROGS pipeline (Galaxy Solutions from Genotoul Bioinformatic platform http://frogs.toulouse.inra.fr) (Escudié *et al*., 2018). Briefly, dereplicated sequences were clustered with Swarm (aggregation parameters equal 1), chimera detection and deletion was based on VSEARCH using the de novo UCHIME method, and sequences were then filtered to keep the minimum number of operational taxonomy units (OTUs) below 0.00005 (Mahe *et al*., 2014; Edgar *et al*., 2010; Rognes *et al*., 2016; Bokulich *et al*., 2013). Sequence affiliation was done using a basic local alignment search tool (BLAST) on silva138_ pintail100 16S database for bacterial sequences and Unite_Fungi_8.2_20200204 database for fungal sequences and a sequence aggregation step was then performed using 98 % of identity and a coverage of 98 %. For fungal data *G. trabeum* sequences were removed. The number of sequences per sample was normalized by random sampling to 2,886 and 3,488 respectively for fungi and bacteria (Table S4). Bacterial OTUs belonging to several proteobacteria classes (alpha, beta and gamma) or the Bacteroidetes phylum were assigned as copiotrophs, whereas deltaproteobacteria and acidobacteria were categorized as oligotrophic organisms (Fierer *et al*., 2007). The trophic mode of fungal OTUs was assigned using FUNGuild (Nguyen *et al*., 2016). Fungi were parsed between saprotroph, pathogenic, symbiotrophic respectively sub-divided in soft rot and white rot fungi, in plant or animal pathogens and fungal parasites, and in endophytes, ericoid mycorrhizal and ectomycorrhizal fungi. Another guild was created to regroup all OTU potentially corresponding to “yeast” (Viotti *et al*., 2021).

### Statistical analyses

Statistical analyses and data visualization were performed with R using phyloseq and ggplot2 packages (Mcmurdie & Holmes, 2013). OTU richness and Shannon’s index were determined using a Phyloseq package (https://rdrr.io/bioc/phyloseq/man/phyloseq-package.html), and Pielou’s evenness was then calculated according to the formula present in the vegan package (Oksanen *et al*., 2019). Following this, pairwise comparisons using the Wilcoxon rank sum test were performed to determine differences between environments. Differences in bacterial and fungal OTU composition were visualized with principal coordinates analyses (PCoA) on Unirac distance matrices using the plot_ordination function in the Phyloseq package. PCoA was used rather than principal components analysis to better reflect the 2-D distance between samples. Permutational multivariate analysis of variance (PERMANOVA) was performed on Unifrac matrices using the Adonis function of the vegan package in order to determine the effect of wood on the bacterial and fungal community structures as well as measuring the impact of degradation inside wood environments. Bacterial and fungal co-occurrence was analyzed using the Corrplot package in R by selecting the Spearman correlation and recognizing only significant values (p value < 0.001).

## Acknowledgements

This work was supported by the Laboratory of Excellence Arbre (ANR-11-LABEX-0002-01; BRAWO project). The UMR1136 and the UR1138 are supported by the French Agency through the Laboratory of Excellence Arbre (ANR-11-LABX-0002-01). Goodell was supported by the National Institute of Food and Agriculture, U.S. Department of Agriculture, the Center for Agriculture, Food and the Environment, and the Microbiology department at University of Massachusetts Amherst, under project number S1075 - MAS00503. Alfredsen was supported by the NIBIO strategic project 11077.The contents are solely the responsibility of the authors and do not necessarily represent the official views of the USDA or NIFA. The Breuil-Chenue site belongs to French national research infrastructure ANAEE-France (ANR-11-INBS-0001). We gratefully acknowledge the financial support received successively from GIP ECOFOR, AllEnvi, ANAEE France, the LTSER Zone Atelier Bassin Moselle and INRAE (DISC, ECODIV). We would also like to thank l’Office National des Forêts, the Parc Naturel Régional du Morvan and Serge Didier for his contribution to the Breuil-Chenue site management.

