## Supplementary figures and images for "Impact of Norway spruce pre-degradation stages induced by *G. trabeum* on fungal and bacterial communities"

### Supplemental Figure 1

#### Slide 1
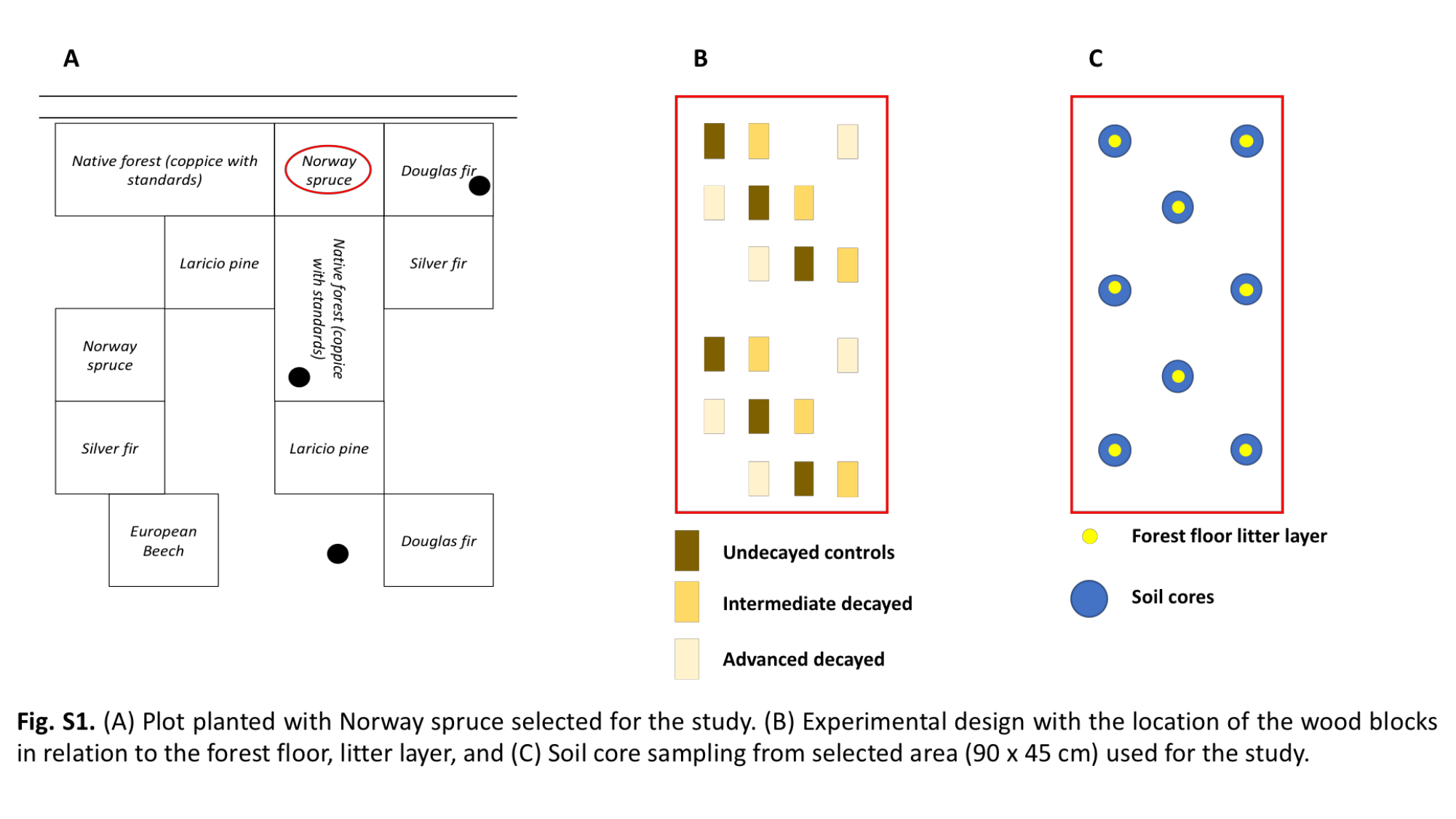
